# Molecular Determinants of Tissue Specificity of Flavivirus Nonstructural Protein 1 Interaction with Endothelial Cells

**DOI:** 10.1101/2022.04.28.489972

**Authors:** Nicholas T.N. Lo, Susan Roodsari, Nicole R. Tin, Scott B. Biering, Eva Harris

## Abstract

Members of the mosquito-borne flavivirus genus such as dengue (DENV), West Nile (WNV), and Zika (ZIKV) viruses cause distinct diseases and affect different tissues. We previously found that the secreted flaviviral nonstructural protein 1 (NS1) interacts with endothelial cells and disrupts endothelial barrier function in a tissue-specific manner consistent with the disease tropism of the respective viruses. However, the underlying molecular mechanism of this tissue-specific NS1-endothelial cell interaction is not well understood. To elucidate the distinct role(s) that the domains of NS1 (β-roll, wing, and β-ladder) play in NS1 interactions with endothelial cells, we constructed flavivirus NS1 chimeras that exchanged the wing and β-ladder domains in a pair-wise manner between DENV, WNV, and ZIKV NS1. We found that both the NS1 wing and β-ladder domains conferred NS1 tissue-specific endothelial dysfunction, with the wing conferring cell binding and the β-ladder involved in inducing endothelial hyperpermeability as measured by trans-endothelial electrical resistance assay. To narrow down the amino acids dictating cell binding specificity, we utilized the DENV-WNV NS1 chimera and identified residues 91 to 93 (GDI) of DENV NS1 as a molecular motif determining binding specificity. Further, using an *in vivo* mouse model of localized leak, we found that the GDI motif of the wing domain was essential for triggering DENV NS1-induced vascular leak in mouse dermis. Taken together, we identify molecular determinants of flavivirus NS1 that confer NS1 binding and vascular leak and highlight the importance of the NS1 wing domain for flavivirus pathogenesis.

**Importance:** Flavivirus NS1 is secreted into the bloodstream from infected cells during a viral infection. Dengue virus NS1 contributes to severe dengue pathology such as endothelial dysfunction and vascular leak independently of the virus. We have shown that multiple flavivirus NS1 proteins result in endothelial dysfunction in a tissue-specific manner consistent with their respective viral tropism. Here, we aimed to identify the molecular determinants that make some, but not other, flavivirus NS1 proteins bind to select endothelial cells in vitro and cause vascular leak in a mouse model. We identified the wing domain of NS1 as a primary determinant conferring differential endothelial dysfunction and vascular leak and narrowed the contributing amino acid residues to a three-residue motif within the wing domain. The insights from this study pave the way for future studies on the effects of flavivirus NS1 on viral dissemination and pathogenesis and offer potential new avenues for antiviral therapies.

## Introduction

Dengue virus (DENV) is a mosquito-borne positive-stranded RNA virus of the *Flavivirus* genus, consisting of four serotypes (DENV1-4). DENV causes ∼100 million symptomatic infections per year globally, ranging from classic dengue fever to the more severe dengue haemorrhagic fever and dengue shock syndrome (1–3). The severe forms of dengue are characterized by clinical presentation of vascular leak as a result of endothelial dysfunction (4). Traditionally, endothelial dysfunction has been attributed to a hyperactivated immune response as a result of uncontrolled viral infection and immune cell activation (5, 6). Recently, we and others have demonstrated that the secreted nonstructural protein 1 (NS1) directly contributes to endothelial dysfunction and vascular leak through interactions with endothelial and immune cells that result in the breakdown of endothelial barriers such as the endothelial glycocalyx and tight and adherens junctions (7–13). In addition to DENV, West Nile (WNV) and Zika (ZIKV) viruses are also members of the *Flavivirus* genus that cause human diseases of public health significance. Like DENV NS1, the NS1 proteins of WNV and ZIKV have also been shown to modulate endothelial barriers. WNV NS1 also has been shown to contribute to WNV infection in the brain (14), while ZIKV NS1 has been shown to modulate the barrier integrity of placental explants as well as the testis (10), contributing to ZIKV infection of Sertoli cells (9, 15).

NS1 is a ∼55-kDa glycoprotein that is highly conserved across the flaviviruses (16). It dimerizes in the endoplasmic reticulum and is found as a component of the viral replication complex. The dimers can also trimerize to form hexamers that are secreted by DENV-infected cells into the bloodstream as a soluble lipoprotein containing lipid cargo (17–19). In severe dengue patients, the levels of detectable NS1 in the blood can be as high as 1-10 µg/mL (20, 21). Secreted NS1 has been shown to associate with components of the innate immune system such as toll-like receptors (13, 22) and complement proteins (23–25), as well as components of the blood-clotting cascade such as platelets (26–28), which results in both host immune evasion and modulation of barrier integrity of endothelial cells. Separately, NS1 can also directly disrupt endothelial cell barrier integrity by promoting the degradation of endothelial glycocalyx components that line the surface of endothelial cells (7, 12, 29, 30) and disrupting junctional proteins that mediate cell-cell interactions (8, 31, 32). In mouse models, NS1 vaccination has been shown to be protective against lethal DENV challenge. In contrast, addition of NS1 to sub-lethal DENV infection was shown to exacerbate pathology resulting in a lethal infection, suggesting the potential of NS1 as a vaccine component and therapeutic target (12).

NS1 proteins from multiple flaviviruses have been shown to interact with endothelial cells of distinct tissue origin in a manner consistent with the tropism of its respective flavivirus (33). For example, DENV NS1 interacts with multiple cell lines including human pulmonary microvascular endothelial cells (HPMEC), which correlates with the systemic nature of DENV infection that can be observed in the lungs. In contrast, WNV NS1 interacts well with brain endothelial cells but minimally with HPMEC, as WNV causes neurological pathologies, such as meningitis and encephalitis, but not pulmonary pathology. Similarly, ZIKV NS1 interacts well with both brain and umbilical vein endothelial cells but minimally with HPMEC, as ZIKV causes neurological and congenital but not pulmonary pathologies. However, how these tissue-specific NS1 interactions are modulated remains unclear.

Flavivirus NS1 contains three domains that are highly conserved: the β-roll (residues 1–29), wing (residues 30–180), and β-ladder (residues 181–352) (34). The NS1 β-roll and wing domains contain many hydrophobic residues that are predicted to interact with the cell’s plasma membrane (35, 36). Interestingly, these surface-exposed residues are conserved among DENV serotypes but divergent across the flavivirus genus (e.g. between DENV and WNV), suggesting their possible roles in mediating NS1 tissue-specific interactions. Additionally, prior studies have suggested that the three domains of NS1 may have distinct functions. DENV NS1 wing domain has been shown to be immunodominant (12, 37, 38), where conserved, hydrophobic residues in the flexible loop (residues 108–129) within the wing domain (Trp-115, Trp-118, Gly-119) have been identified to partially contribute to DENV NS1 binding to HPMEC (35, 39). In contrast, select residues in the β-ladder domain (such as residues N207, A303, E326, and E327) have been implicated as important for NS1-induced endothelial hyperpermeability, while being dispensable for binding endothelial cells (31, 39). While these data offer clues about NS1 molecular determinants of tissue-specific interactions with endothelial cells, a systematic investigation of these different domains in distinct flavivirus NS1 proteins has not been undertaken.

In this study, we provide new insights on the tissue-specific interactions between flavivirus NS1 and endothelial cells using NS1 chimeras that exchange the wing and β-ladder domains of DENV, WNV, and ZIKV in a pair-wise manner with one another. We found that both the wing and β-ladder domains confer tissue-specific barrier dysfunction, with the wing influencing initial attachment to endothelial cells and the β-ladder involved in inducing endothelial hyperpermeability *in vitro*. We further identified a variable 3-amino acid (aa) motif in the wing domain as a molecular determinant for tissue specificity. Finally, we identify the wing domain and the 3-aa motif as a driver of DENV-triggered vascular leak *in vivo*. Taken together, our results provide insights into the tissue-specific interactions of flavivirus NS1.

## Results

### The wing domain of flavivirus NS1 confers binding to endothelial cells

In previous work, we showed that DENV NS1 binds to human lung endothelial cells (HPMEC) at higher levels than WNV and ZIKV NS1 (33), correlating with the capacity of DENV NS1 but not WNV or ZIKV NS1 to cause endothelial hyperpermeability of HPMEC and pleural effusion in humans. Similarly, DENV and ZIKV NS1 exhibited higher binding than WNV NS1 to human umbilical vein microvascular endothelial cells (HUVEC) and caused endothelial hyperpermeability of HUVEC (33). To uncover the molecular determinants of flavivirus NS1 that dictate this tissue-specific endothelial cell tropism, we generated chimeric NS1 proteins that exchanged either the wing or the β-ladder domains between DENV and WNV, WNV and ZIKV, and DENV and ZIKV NS1, respectively (Fig. 1A, D, G). We cloned and expressed the C-terminally His-tagged NS1 proteins in HEK-293 cells, which were successfully secreted, and then purified NS1 oligomers using cobalt affinity chromatography (Fig. 2). Chimeric proteins are annotated by their 3 domains from the respective flavivirus NS1 proteins in sequence “β-roll–wing–β-ladder” (e.g., D-D-W designating DENV β-roll – DENV wing – WNV β-ladder). We treated HPMEC with either wild-type (WT) or DENV-WNV chimeric NS1 proteins and measured the levels of NS1 binding to HPMEC using an immunofluorescence microscopy assay (IFA). We found that the chimeric NS1 constructs containing DENV NS1 wing domain bound to HPMEC at comparable levels to WT DENV NS1, whereas the constructs containing WNV NS1 wing exhibited significantly lower binding than WT DENV NS1, but at comparable levels to WT WNV NS1 (Fig. 1B, C). A similar pattern was observed when we treated HUVEC with WNV-ZIKV chimeric NS1, where constructs that contained ZIKV NS1 wing bound to HUVEC at comparable levels to WT ZIKV NS1, whereas the constructs containing WNV NS1 wing bound to HUVEC at lower levels consistent with WT WNV NS1 (Fig. 1E, 1F). We expanded this chimeric NS1 approach to include DENV and ZIKV NS1 and found that while constructs with ZIKV NS1 wing significantly diminished binding to HPMEC, the constructs with DENV NS1 wing did not gain binding to HPMEC comparably to WT DENV NS1 (Fig. 1H, J), possibly due to differential effects of interdomain interactions between flaviviruses. Overall, these results indicate that the tissue-specific patterns of flavivirus NS1 binding to endothelial cells are driven by the wing domain of NS1.

**Figure 1.**
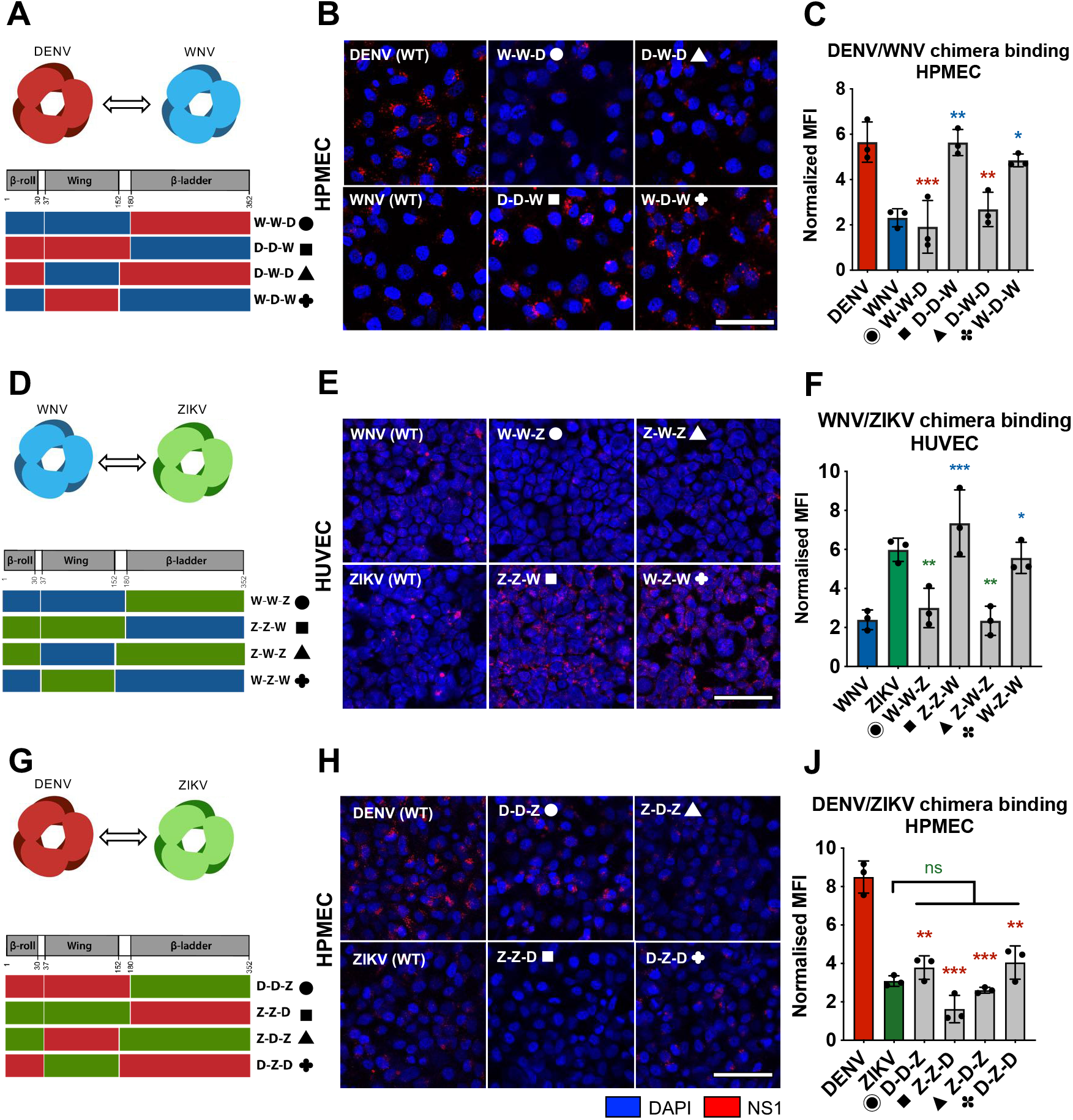
The wing domain of flavivirus NS1 confers binding to endothelial cells. **(A)** Schematic representation of chimeric NS1 proteins that exchange wing or β-ladder domains between DENV and WNV NS1. Red box represents DENV NS1, and blue box represents WNV NS1. The notation (e.g., D-D-W) indicates the unique flavivirus NS1 domain at β-roll, wing, β-ladder domains. **(B)** Recombinant WT or chimeric NS1 (10 µg/mL) was added to HPMEC and incubated at 37°C for 1h. NS1 binding was assessed by immunofluorescence microscopy, and representative images from three experiments are shown. **(C)** Quantification of (B), normalized to untreated controls. **(D, G)** Schematic representation of chimeric NS1 produced with WNV and ZIKV NS1 (D) or DENV and ZIKV NS1 (G), respectively. Green box represents ZIKV NS1. **(E, H)** NS1 binding to HUVEC or HPMEC, as in B. **(F, J)** Quantification of (E) and (H), respectively. MFI, mean fluorescence intensity. Images represent n=3 biological replicates. Scale bars are 100µm. Data plotted as mean ± SEM. *p<0.05, **p<0.01, ***p<0.005, ****p<0.001 by one-way ANOVA with multiple comparisons. Star colors indicate the respective control each construct was compared to.

**Figure 2.**
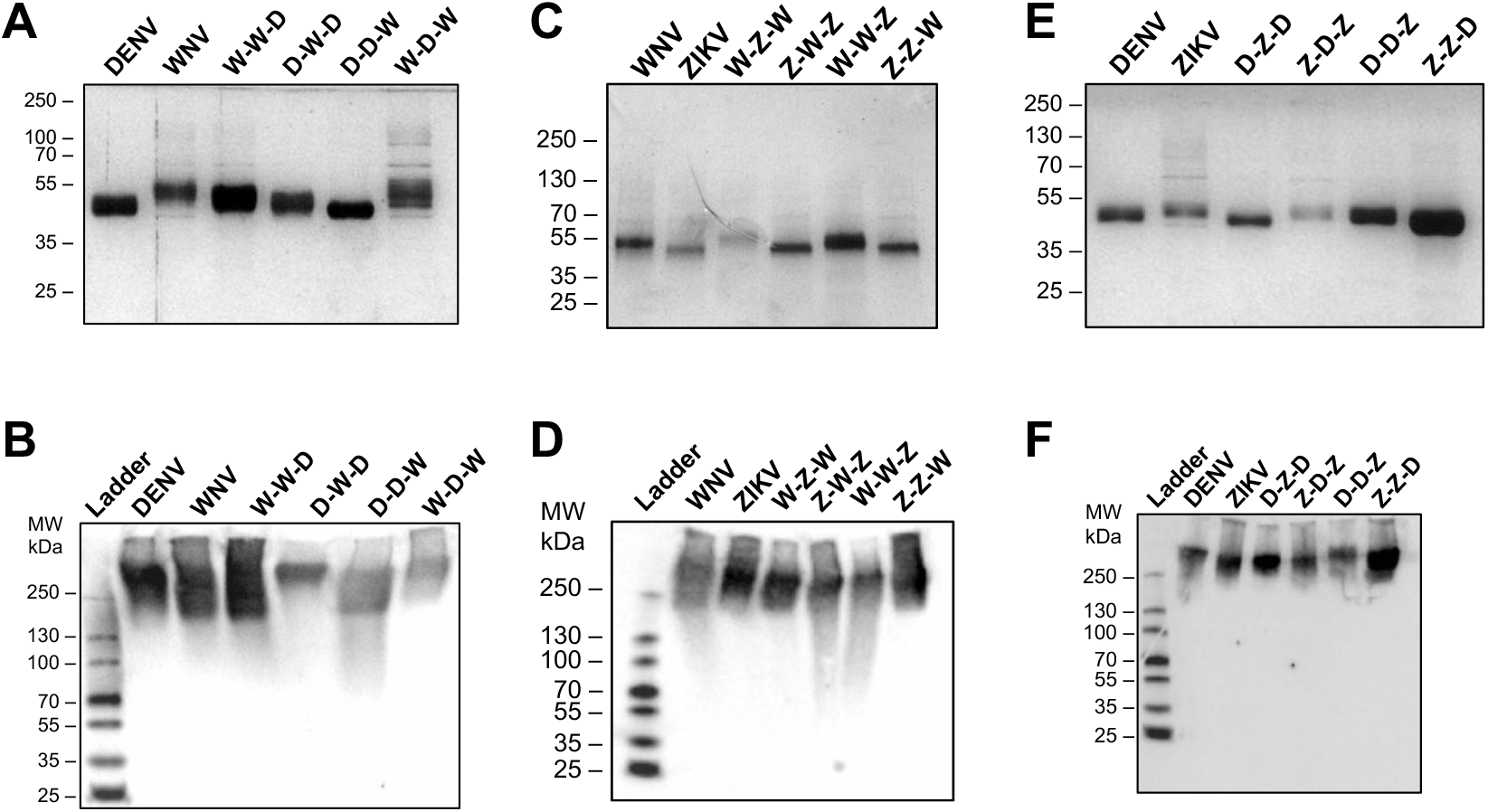
Production and quality control of flavivirus NS1 chimeric proteins. Proteins were evaluated by silver stain following SDS-PAGE (A, C, E) and native PAGE (B, D, F). 2µg of each NS1 construct was used. For native gels, proteins were detected using anti-His mAb. **(A)** Silver stain of DENV-WNV NS1 chimeras. **(B)** Native PAGE of DENV-WNV NS1 chimeras. **(C)** Silver stain of WNV-ZIKV NS1 chimeras. **(D)** Native PAGE of WNV-ZIKV NS1 chimeras. **(E)** Silver stain of DENV-ZIKV NS1 chimeras. **(F)** Native PAGE of DENV-ZIKV NS1 chimeras.

### The wing and β-ladder domains of flavivirus NS1 proteins are necessary for inducing endothelial hyperpermeability

Following binding to endothelial cells, DENV NS1 is taken up by cells via clathrin-mediated endocytosis, trafficks to endosomes, and activates enzymes that degrade the endothelial glycocalyx and disrupt intercellular junctional complexes (7, 10, 31). This results in disruption of endothelial barrier integrity, making endothelial cells hyperpermeable to solutes and liquids. We have previously shown that DENV NS1 can cause endothelial barrier dysfunction by inducing endothelial hyperpermeability independently of the virus (7, 30). Given this finding, we next explored whether the NS1 wing domain that conferred DENV NS1 binding to HPMEC is also responsible for causing HPMEC hyperpermeability. To investigate how different flavivirus NS1 domains influence the capacity of NS1 to trigger endothelial hyperpermeability, we utilized the DENV-WNV NS1 chimeras in a trans-endothelial electrical resistance (TEER) assay. We treated HPMEC seeded in the apical chamber of a trans-well with either WT or chimeric NS1 (2.5 μg/ml) and measured the TEER values between the apical and the basolateral chambers of the transwell (Fig. 3A, B). Consistent with previous observations (33), WT DENV NS1 resulted in significantly greater TEER reduction of HPMEC than WNV NS1, indicating greater endothelial barrier hyperpermeability. Interestingly, the constructs with either the DENV wing or β-ladder domain (i.e. all of the mutant NS1 chimeras) were unable to cause TEER reduction to the same extent as WT DENV NS1, indicating that while the wing domain confers cell binding, both the DENV wing and the DENV β-ladder domains are required for NS1 to trigger endothelial hyperpermeability of HPMEC.

**Figure 3.**
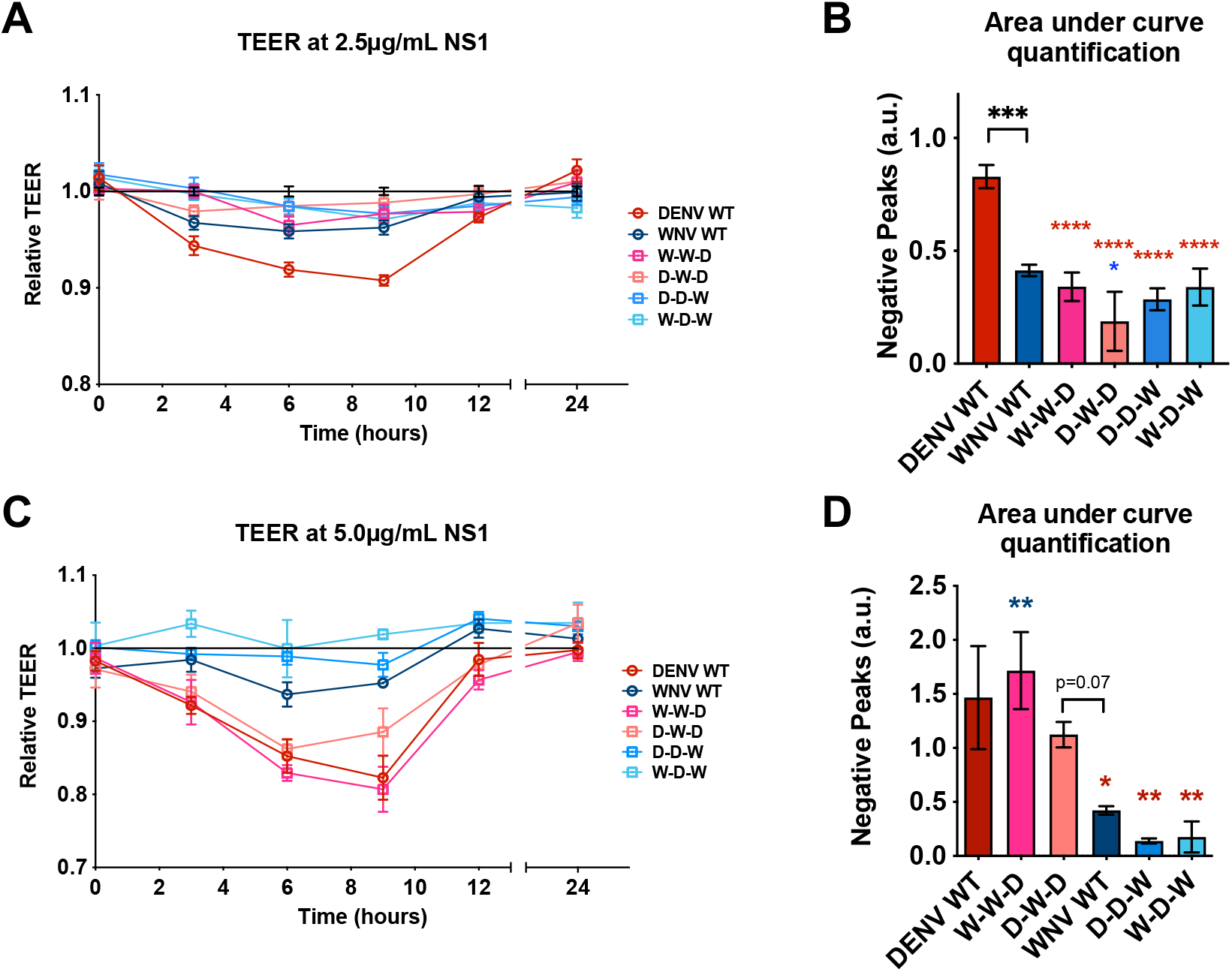
The wing and β-ladder domains of flavivirus NS1 are necessary for inducing endothelial cell hyperpermeability. **(A)** HPMEC monolayer was seeded in the apical chambers of 24-well trans-well inserts and treated with either WT or chimeric NS1 proteins at 2.5µg/mL. The trans-endothelial electrical resistance (TEER) between the apical and basolateral chamber were measured over time and normalized to the untreated controls at each respective timepoint. **(B)** Quantification of the area between the curve and Y=1.0 in (A) (“area under the curve”), correlating to a decrease in electrical resistance. **(C)** Endothelial permeability assay as in (A), but with the indicated NS1 protein at 5µg/mL. **(D)** Area under the curve of (C). A.u., arbitrary units. Data represent at least n=3 biological replicates, plotted as mean ± SEM. *p<0.05, **p<0.01, ***p<0.005, ****p<0.001 by one-way ANOVA with multiple comparisons. Star colors indicate the respective control each construct was compared to.

Since we have previously observed that NS1 triggers endothelial hyperpermeability in a dose-dependent manner (7, 8), we tested whether our chimeric NS1 proteins would trigger endothelial hyperpermeability when cells were treated with a higher concentration. We repeated the TEER experiments using 5 µg/mL of our NS1 proteins (Fig. 3C, D), and observed that the constructs containing the DENV NS1 β-ladder (W-W-**D** and D-W-**D**) caused a decrease in TEER on HPMEC comparably to WT DENV NS1, whereas the constructs containing the WNV β-ladder domain (D-D-**W** and W-D-**W**) did not cause endothelial hyperpermeability, similar to WT WNV NS1. This implicates a role for the DENV β-ladder in inducing endothelial hyperpermeability as measured by TEER, consistent with our previous findings (31, 39).

### A 3-amino acid motif in the wing domain of DENV NS1 confers endothelial cell binding specificity between DENV and WNV NS1

While NS1 is highly conserved across the flavivirus genus (40), areas of variability can be found within the wing domain, specifically between amino acid (aa) residues 90 and 120 (Fig. 4A). We have previously identified the same stretch of residues within the NS1 wing domain to be immunodominant, eliciting robust antibody responses in DENV2-infected mice and natural human infections (37). Multiple structural studies have reported a flexible loop structure located near residues 90-120 that is thought to be critical for endothelial cell binding, given its predicted proximity to cell membranes and its exposure on the surface of dimeric NS1 (41–43). This suggests a functional importance that could explain the tissue-specificity of flavivirus NS1. As such, we hypothesized that amino acids within residues 90 to 120 of the wing domain confer tissue-specific endothelial binding.

**Figure 4.**
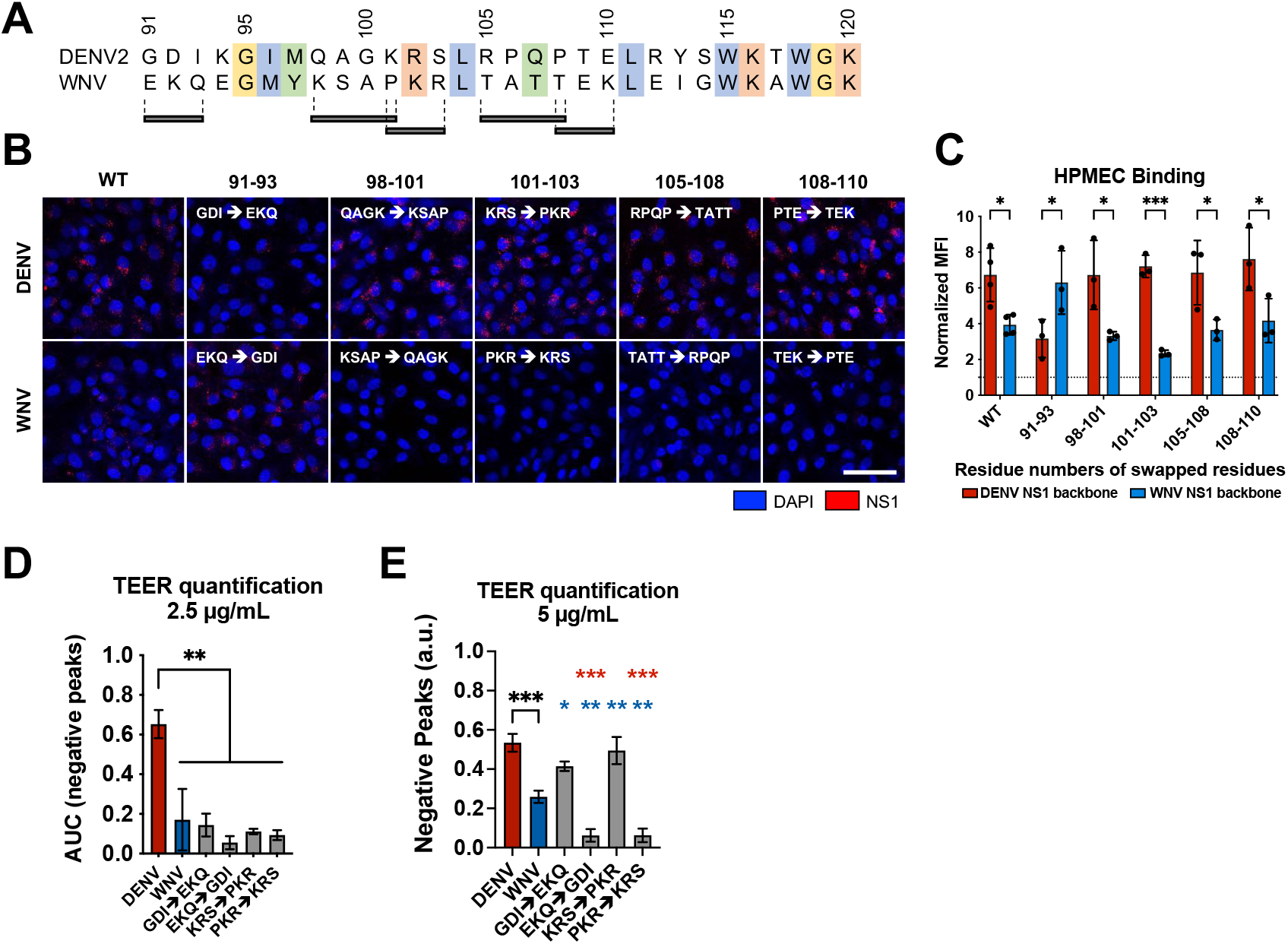
Residues 91-93 of DENV NS1 drive DENV NS1-endothelial cell interactions. **(A)** Sequence alignment of DENV and WNV NS1 from residue 91 to 120, with black bars indicating the 3-4 residue motifs swapped in the site-directed mutants between DENV and WNV NS1. Colors of the residues indicate their similarities based on biochemical properties. **(B)** NS1 binding assay where 10µg/mL of WT or mutant NS1 as indicated were added to HPMEC and imaged using immunofluorescence microscopy. **(C)** Quantification of (B). **(D)** HPMEC monolayer was seeded in the apical chamber of a trans-well and treated with either WT or site-directed mutant NS1 proteins at 2.5µg/mL. Trans-endothelial electrical resistance (TEER) was measured over time, normalized to the untreated controls of respective timepoints. Area-under-curve quantification TEER curves is shown. **(E)** Same as (D) but treating with NS1 proteins at 5.0 µg/mL. Area-under-curve quantification of TEER curves is shown. All data in this figure are from at least 3 biological replicates, plotted as mean ± SEM. *p<0.05, **p<0.01, ***p<0.005, ****p<0.001 by one-way ANOVA with multiple comparisons. a.u., arbitrary units. Star colors indicate the respective control each construct has been compared to.

To test this hypothesis, we aligned the DENV and WNV NS1 sequences to identify differentially conserved residues that were predicted to be surface-exposed, reasoning that these divergent residues may dictate tissue specificity of NS1 (42, 43). We identified five sites of 3-4aa between residues 90 and 120 that contained divergent residues across flaviviruses, which were conserved within each flavivirus (Fig. 4A). To test the involvement of these residues in the differential DENV-WNV NS1 cell binding phenotype, we generated 5 pairs of site-specific mutants that exchanged these motifs between DENV and WNV NS1 (Figs. 4 and 5). We then treated HPMEC with either WT NS1 or site-specific NS1 mutants and measured the levels of NS1 binding to HPMEC by IFA (Fig. 4B). Interestingly, we found that while 4 pairs retained the binding pattern of their parental NS1 protein, one pair exhibited the opposite pattern. WNV NS1 containing DENV residues 91-93 (WNV-GDI) gained the capacity to bind to HPMEC at comparable levels to WT DENV NS1, whereas the DENV NS1 containing residues 91-93 from WNV NS1 (DENV-EKQ) had reduced capacity to bind to HPMEC, exhibiting levels similar to WT WNV NS1 (Fig. 4B, C). These results suggest that residues 91-93 of DENV NS1 form a motif that contributes to the DENV-WNV tissue-specific endothelial cell binding pattern we observe.

**Figure 5.**
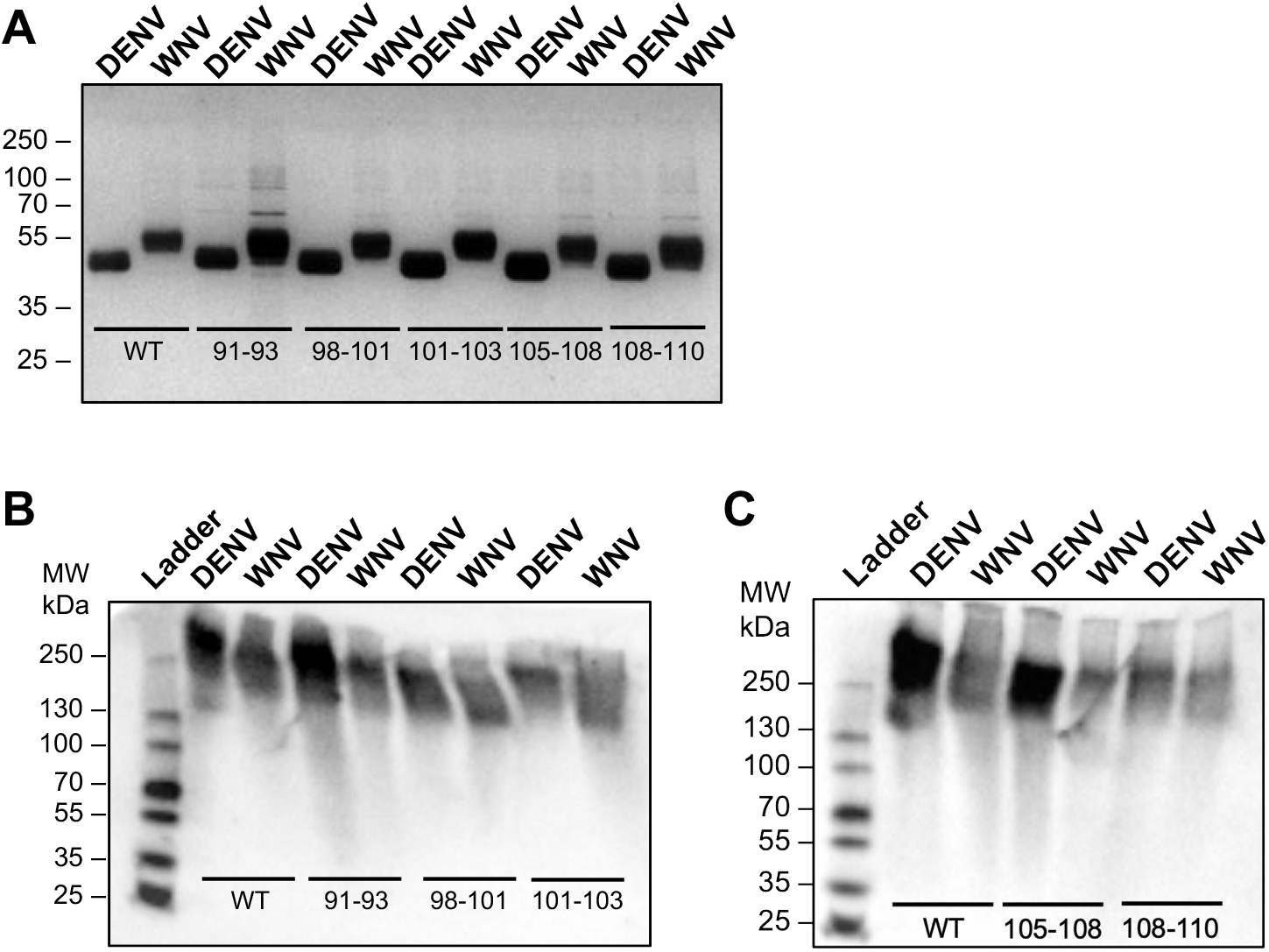
Production and quality control of flavivirus NS1 site-directed mutants. Proteins were evaluated by silver stain following SDS-PAGE gel (A) and native PAGE (B, C). 2µg of each NS1 construct was used. For native gels, proteins were detected using anti-His mAb. Top row labels indicate the flavivirus NS1 backbone. Numbering under bands refer to the residues that were swapped between DENV and WNV NS1. **(A)** Silver stain of site-directed NS1 mutants between DENV and WNV NS1. **(B, C)** Native PAGE of site-directed NS1 mutants between DENV and WNV NS1, split into two gels each with respective WT DENV and WNV NS1 controls. WT, wild-type.

To determine if residues 91-93 conferring NS1 cell binding specificity between DENV/WNV also dictate the capacity of DENV/WNV NS1 to trigger endothelial hyperpermeability, we conducted a TEER assay on HPMEC with NS1 mutants swapped at residues 91-93 and 101-103 for comparison (Fig. 4D). Similar to our observations above, at low NS1 levels (2.5 µg/mL), only WT DENV NS1 was able to induce endothelial hyperpermeability, in contrast to WT WNV and mutants from the DENV/WNV 91-93 and 101-103 swaps that did not noticeably alter permeability, suggesting that while the residues in the wing domain are important for inducing endothelial hyperpermeability, they alone are not sufficient.

We next repeated the TEER experiments at 5µg/mL to confirm if the wing deficiency phenotype of DENV NS1 could be overcome by utilization of a higher dose of NS1 (Fig. 4E). WT DENV NS1 and NS1 constructs containing WNV motifs of either residues 91-93 and 101-103 (both containing DENV β-ladder) were able to trigger endothelial hyperpermeability, whereas WT WNV NS1 caused less TEER decrease, and WNV NS1 constructs containing DENV NS1 motifs of both residues 91-93 and 101-103 (both containing WNV β-ladder) remained unable to cause TEER decrease. Together, these results are consistent with the previous TEER data involving NS1 domain chimeras (Fig. 3), where at the higher NS1 concentration of 5 µg/mL, the constructs containing DENV β-ladder triggered endothelial hyperpermeability.

### The wing domain of DENV NS1 confers NS1-induced vascular leak *in vivo*

To explore if the tissue-specific NS1-endothelial cell interactions we observed *in vitro* could be recapitulated *in vivo*, we next asked which NS1 domains were required for tissue-specific vascular leak *in vivo*. We used a mouse model of localized dermal leak, in which we have previously shown that DENV NS1 causes significantly higher leak than WNV NS1 (30). We shaved the hair from the back (dorsal side) of wild-type C57BL/6 mice and administered injections intradermally (ID): PBS as baseline vehicle control, WT NS1, and DENV-WNV NS1 chimeras. Immediately following ID injections, we retro-orbitally administered dextran conjugated to Alexa-680. Two hours post-NS1 treatment, we excised the dorsal dermis and used a fluorescent scanner to measure the extent of vascular leak as indicated by dextran-associated fluorescence, since dextran is a small molecule that can extravasate into tissues (Fig. 6A, B).

**Figure 6.**
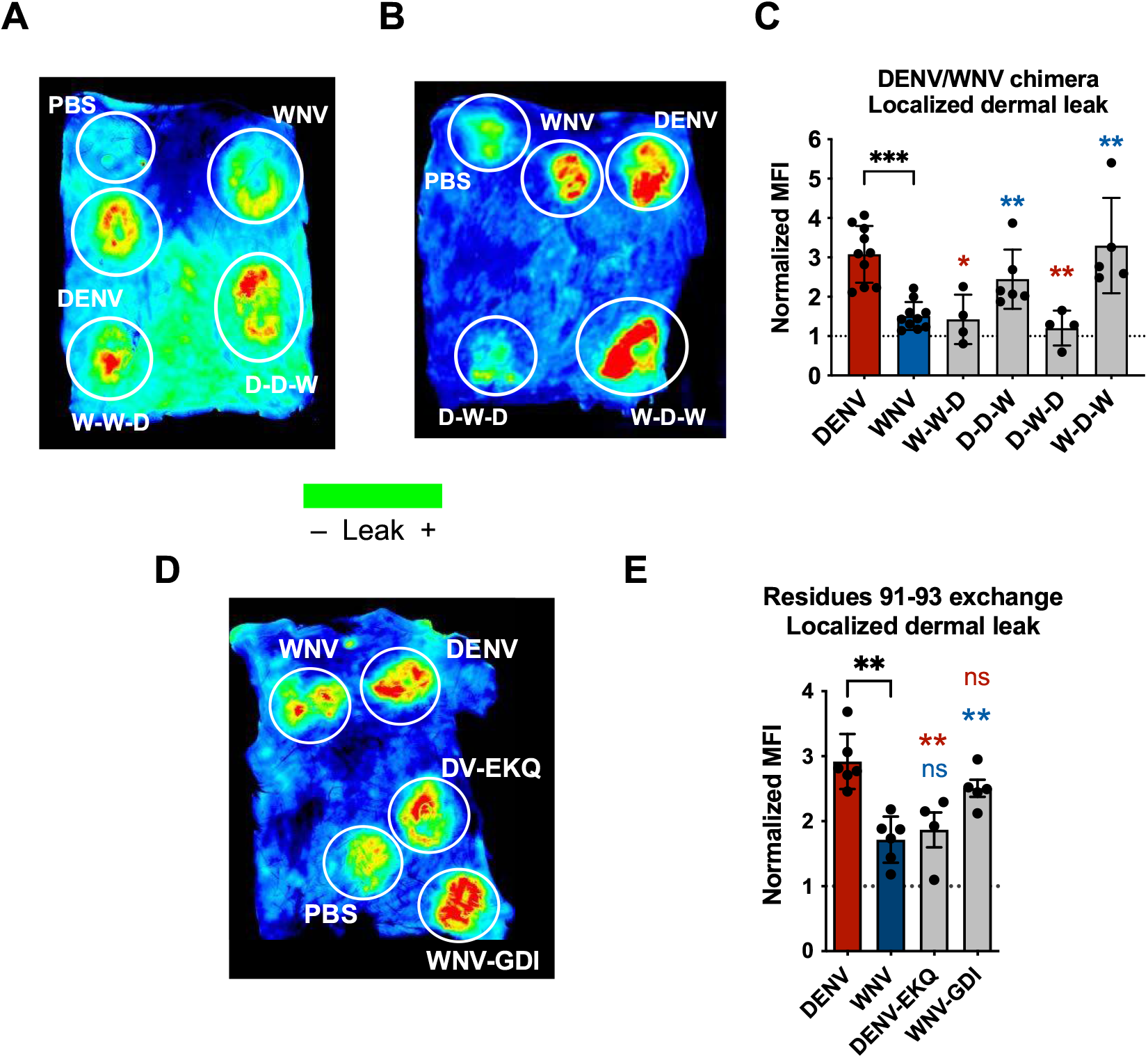
The wing domain of DENV NS1 and residues 91 to 93 within the wing domain confer NS1-induced vascular leak *in vivo*. **(A, B)** Wild-type C57BL/6J mice were shaved by removing hair from their backs 3 days before experiment. On the day of experiment, PBS, WT and chimeric NS1 proteins (15 µg) were administered intradermally into discrete spots on the shaved dorsal dermis. Immediately following NS1 injections, 25 µg of Dextran conjugated to Alexa Fluor 680 was administered retro-orbitally. Two hours post-injection, the dermis of each mouse was collected and visualized using a fluorescent scanner. Representative images of 4 to 6 mice backs are shown. Chimeric proteins were analyzed in different mice with each mouse containing both positive and negative controls (WT DENV and WNV NS1, respectively). **(C)** Mean fluorescent intensity (MFI) quantification of (A) and (B), normalized to PBS injection. **(D)** Same as (A) and (B), but using site-directed NS1 mutants that exchanged residues 91 to 93 between DENV and WNV NS1. Representative image of 4 mice backs are shown. **(E)** MFI quantification of (D), normalized to PBS injection. Data plotted as mean ± SEM. *p<0.05, **p<0.01, ***p<0.005, ****p<0.001 by unpaired Mann-Whitney U test. Star colours indicate the respective control each construct was compared to.

Using this model, we found that the NS1 chimeras containing DENV wing domain (D-**D**-W and W-**D**-W) caused comparable levels of leak as WT DENV NS1, which is significantly higher leak than the leak caused by WT WNV NS1 (Fig. 6C). Conversely, the NS1 chimeras containing WNV wing domain – and DENV β-ladder (W-**W**-D and D-**W**-D) – caused leak comparable to WT WNV NS1, and significantly less than WT DENV NS1. These data suggest that the DENV NS1 wing is driving NS1-induced vascular leak in the dorsal dermis of mice *in vivo*.

Since we identified residues 91-93 of DENV as a determinant of wing-mediated endothelial hyperpermeability *in vitro*, we asked whether this motif was also a determinant for wing-mediated dermal leak *in vivo*. Using the same dermal leak model as above, we intradermally injected PBS, WT NS1 proteins, and DENV NS1 with WNV NS1 residues 91-93 and *vice versa*, followed by retro-orbital administration of Dextran-A680 (Fig. 6D). We found that WNV NS1 with DENV residues 91-93 gained the ability to trigger vascular leak at comparable levels as WT DENV NS1, while DENV NS1 with WNV residues 91-93 caused significantly less leak, comparable to WT WNV NS1 (Fig. 6E). Taken together, these results suggest that the DENV NS1 wing domain, along with residues 91-93 (GDI), are required to trigger localized vascular leak in this *in vivo* model.

## Discussion

In this study, we determined that both the wing and β-ladder domains of flavivirus NS1 proteins drive differential interactions with human endothelial cells, with the wing influencing binding to endothelial cells and the β-ladder involved in inducing endothelial hyperpermeability *in vitro*. We further identified residues 91 to 93 (GDI) within the flexible loop of the DENV NS1 wing domain as a critical molecular determinant for tissue-specific NS1-endothelial cell binding. Finally, we showed that the wing domain, along with the GDI motif, also dictate the capacity of NS1 to trigger vascular leak *in vivo*. Our finding that the NS1 wing domain contributes to endothelial binding and hyperpermeability is consistent with our previous observations (39) and with studies reporting that in DENV NS1-vaccinated mice, anti-wing antibodies were highly protective against lethal DENV infection (37, 44). While in its cell-bound dimeric form, residues within the 90 to 120 region of the wing domain are expected to interact with the plasma membrane (34, 36, 42, 43), which our findings support. In addition, the hydrophobic residues in the DENV NS1 wing domain have also been implicated to be involved in membrane remodeling (45).

It could be expected that flavivirus NS1 contains both conserved core residues mediating interactions with endothelial cells, as well as divergent residues that confer tissue-specificity. In agreement with this hypothesis, previous work from our group uncovered NS1 residues involved in endothelial cell binding that are conserved across the *Flavivirus* genus (W115-W118-G119); these residues are located in the flexible loop within the wing domain (39). In that study, we mutated the W-W-G residues to alanines; the DENV NS1-WWG mutant bound to HPMEC at lower levels than WT DENV NS1 and had reduced capacity to cause endothelial hyperpermeability in HPMEC. This finding suggests that the highly conserved WWG residues may mediate a baseline level of NS1-endothelial cell binding across all flavivirus NS1 proteins, while the tissue-specificity is likely mediated by non-conserved molecular determinants. The conserved WWG residues contrast the DENV-GDI motif we identified in the present study, which is highly variable among flaviviruses.

One interesting observation in our data was how the DENV-WNV chimeras that bound weakly at 2.5µg/mL to HPMEC (W-**W**-D and D-**W**-D, containing WNV wing domain), at much lower levels than WT DENV NS1, could induce HPMEC hyperpermeability at comparable levels as WT DENV NS1 at the higher NS1 concentration of 5µg/mL (Figure 2). We observed the same trend again when we tested the capacity of the site-directed mutants to cause endothelial hyperpermeability, where at higher concentrations, the DENV-EKQ mutant NS1 bound at a low level to HPMEC while remaining able to cause HPMEC hyperpermeability, whereas WNV-GDI NS1 bound HPMEC but could not cause HPMEC hyperpermeability. This implicates the β-ladder in conferring endothelial cell hyperpermeability, and is consistent with prior studies that identified residues in the β-ladder as important for endothelial hyperpermeability at a step following endothelial cell binding. A glutamine point mutant of the glycosylated residue N207 (31) and targeted mutation at residues T301, A303, E326, E327 (39), which are in the β-ladder domain, were able to bind to HPMEC, but deficient in causing HPMEC hyperpermeability.

It is important to note that while WT WNV NS1 bound to HPMEC less efficiently than DENV NS1, it did bind at detectable levels, although this level of binding was not sufficient to trigger endothelial hyperpermeability (Fig. 3) (14). In the TEER model used to measure endothelial hyperpermeability *in vitro*, cells are treated with NS1 and left undisturbed over time under static conditions, in contrast to DENV infection *in vivo*, where NS1 travels through the bloodstream with constant blood flow. As a result, when the WNV NS1 and DENV-WNV NS1 chimeras (containing WNV wing but DENV β-ladder) that bound poorly to HPMEC were left undisturbed on HPMEC in TEER assays, we hypothesize that their low-level binding was then sufficient to induce hyperpermeability, which was driven by the DENV β-ladder domain. In the *in vivo* environment, we observed that the vascular leak phenotype was conferred by the wing domain, which is consistent with the observation of wing-driven endothelial binding *in vitro*. This suggests that in the fluid condition of the *in vivo* environment, it is critical for NS1 to have strong interactions with host factors on the cell surface such that NS1 binding, which is mediated by the wing domain, can occur despite blood flow.

Further, the complexity of the *in vivo* condition likely accounts for the discrepancy we observe between the *in vitro* endothelial permeability system and the *in vivo* murine dermal leak model, where constructs containing DENV wing and WNV β-ladder could cause vascular leak *in vivo* despite not causing endothelial hyperpermeability *in vitro*. The *in vitro* TEER assay contains only endothelial cells, whereas in the *in vivo* mouse model, other non-endothelial intrinsic factors may be at play, which could be mediated by the wing domain of NS1. In addition, the kinetics differed in our *in vivo* model of localized leak versus *in vitro* TEER assay. Finally, the NS1 proteins might have different effects on the dermal cells in the *in vivo* system as compared to pulmonary cells in the *in vitro* system. Future studies are needed to fully characterize the relative contribution of non-endothelial factors to NS1-mediated vascular leak.

A critical question to address next is the mechanism by which the aa 91-93 motif of the wing domain mediates tissue-specific endothelial cell binding. We propose two ways in which residues 91 to 93 may mediate tissue-specific interactions. One possibility is that the motif directly interacts with specific host factors on the surface of endothelial cells such as glycans or proteins. In particular, the carboxylate side chain of D92 points directly towards the cell surface; it may interact with host factors, distinct from the WWG motif, which may facilitate association with the plasma membrane (Fig. 7). A second possibility is that the aa 91-93 motif modulates the flexibility of the flexible loop containing the WWG motif (39), which is ultimately the site predicted to interact with endothelial cells.

**Figure 7.**
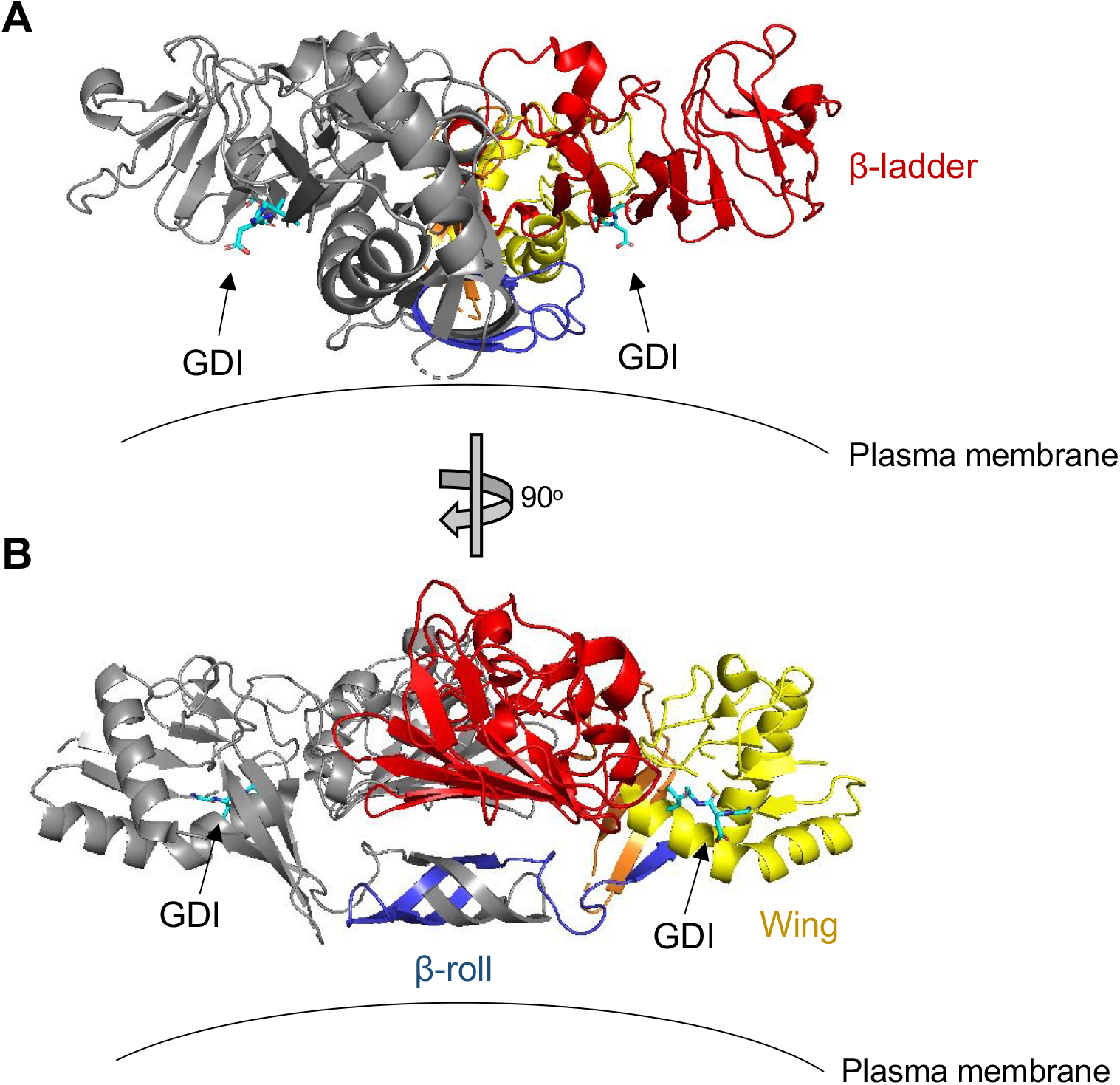
DENV NS1 structure depicting location of aa 91-93 GDI motif. Dimeric DENV2 NS1 structure at 2.89Å (PDB 7K93; Biering*, Akey* *et al*, 2021 [39]) was annotated to show the location of the aa 91-93 GDI motif within the wing domain in spatial and structural reference to the rest of NS1. One monomer is coloured grey while the other monomer is coloured as follows: blue for β-roll, yellow for wing, red for β-ladder, and orange for inter-domain connecting regions. Arrows point towards GDI motif on both monomers, coloured in cyan. Plasma membrane is shown to indicate the position in which NS1 is proposed to interact with. **(A)** and **(B)** are rotated 90° along the Y-axis from each other.

On the other side of NS1 tissue-specific interactions lies the host endothelial cell. The endothelial cell surface is made up of the endothelial glycocalyx, which is a network of luminal membrane-bound glycoproteins and proteoglycans with both short, branched carbohydrates and long, unbranched glycosaminoglycan side-chains (47, 48). Glycocalyx components include sialic acid, heparan sulfate, chondroitin sulfate, and hyaluronan (47). The composition and ratio of the components that make up the glycocalyx can vary significantly depending on the tissue of origin of endothelial cells, which in turn can result in differential binding of viral proteins to distinct tissues. As such, different residues in both the aa 91-93 motif and the broader wing domain may influence the interactions with specific types of glycans on different endothelial cells.

Our study begins to uncover the molecular determinants for flavivirus NS1 binding and tissue tropism. We establish the flavivirus NS1 wing domain to be important for binding endothelial cells and causing vascular leak, and in the case of DENV NS1, identify residues 91 to 93 within the wing domain as key determinants driving such interactions. This new molecular insight into what determines flavivirus NS1 tissue specificity is crucial for understanding pan-flaviviral pathogenesis and offers new approaches for antiviral therapies.

## Acknowledgements

We thank Dr. Janet Smith at the University of Michigan for her helpful discussions about NS1 structure-function relationship. We also thank Evan Juan, Bryan Castillo-Rojas, and Richard Ruan of the Harris laboratory for their technical assistance. Confocal imaging experiments were conducted on a Zeiss LSM 710 microscope at the CRL Molecular Imaging Center at UC Berkeley, which is supported by the Gordon and Betty Moore Foundation. This study was supported by NIH grant R01 AI124493. S.B.B was supported by the Open Philanthropy Life Science Research Foundation postdoctoral award.

N.L., S.B.B, and E.H. conceived the study. N.L., S.R. and N.T. cloned chimeric NS1 constructs. N.L. and S.R. produced and purified the recombinant NS1 proteins. N.L. performed the experiments and data analysis in this study. S.R. assisted on imaging and image analysis. S.B.B. and E.H. provided project guidance. E.H. acquired funding and provided resources. N.L. wrote the initial manuscript draft. N.L., S.B.B, and E.H. reviewed and edited the manuscript and all authors provided editorial comments.

The authors declare no competing interests.

## Materials and Methods

### Cell lines

FreeStyle 293F suspension cells (Thermo Fisher Scientific) were used for production of recombinant NS1 proteins. 293F cells were cultured in FreeStyle 293 Expression medium (Thermo Fisher Scientific) containing 1% penicillin/streptomycin (P/S) and grown in a CO_2_ incubator at 37°C with 8% CO_2_ and maintained on a cell shaker at ∼130 rpm. HPMEC (HPMEC-ST1.6r) were kindly donated by Dr. J.C. Kirkpatrick at Johannes Gutenberg University, Germany, and were used for NS1 cell binding and TEER assays. HUVEC were kindly gifted from Dr. Melissa Lodoen at the University of California, Irvine. HUVEC are primary endothelial cells obtained from a single female donor (Lonza). Both HPMEC and HUVEC cell lines were propagated (passages 5–10) and maintained in endothelial growth medium 2 (EGM-2) using the EGM-2 bullet kit from Lonza following the manufacturer’s specifications and grown in a CO_2_ incubator at 37°C with 5% CO_2_.

### NS1 mutagenesis and cloning chimeric NS1 proteins

Chimeric NS1 proteins were produced by amplifying fragments of β-roll, wing, and β-ladder domains from the WT DENV2 NS1 (Thailand/16681), WNV NS1 (NY99), or ZIKV NS1 (Uganda MR766), using primers listed in Table 1. The N-terminus of β-roll and C-terminus of β-ladder primer sequences were flanked with nucleotide bases complementary to the protein expression vector plasmid “pMAB”. The pMAB vector encodes a N-terminal CD33 signal sequence and C-terminal 6xHis tag, a kind gift from Dr. Michael Diamond, Washington University at St. Louis. The domain fragments and pMAB vector were fused together using overlap extension PCR. Site-directed NS1 mutants were produced using a site-directed mutagenesis kit (QuikChange XL Site-Directed Mutagenesis Kit, Agilent) following the manufacturer’s instructions, with primers listed in Table 1. All mutant NS1 constructs were sequence-verified with 5’ and 3’ primers that recognize the pMAB vector beyond the mutagenesis insertion region.

**TABLE 1.**
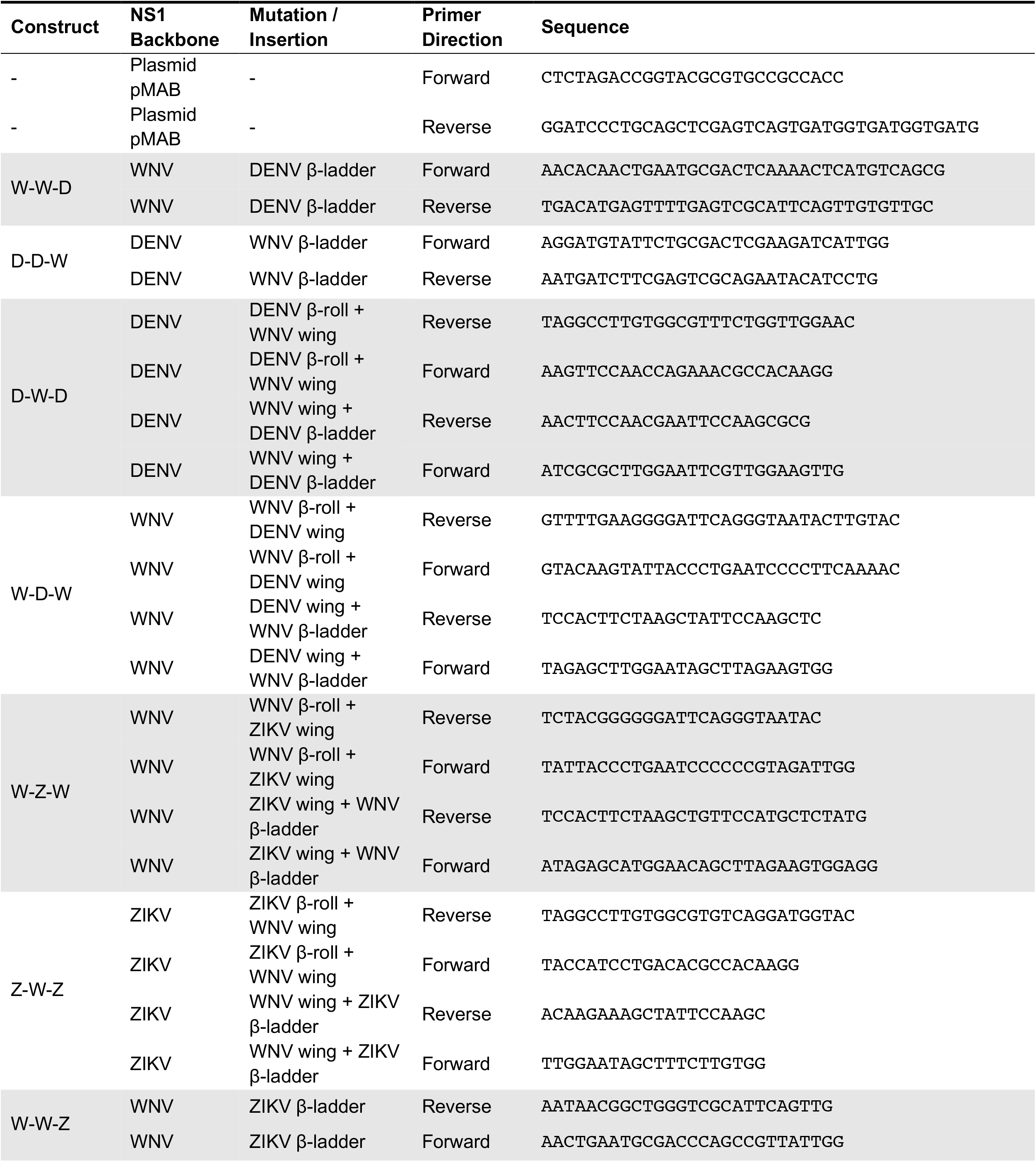
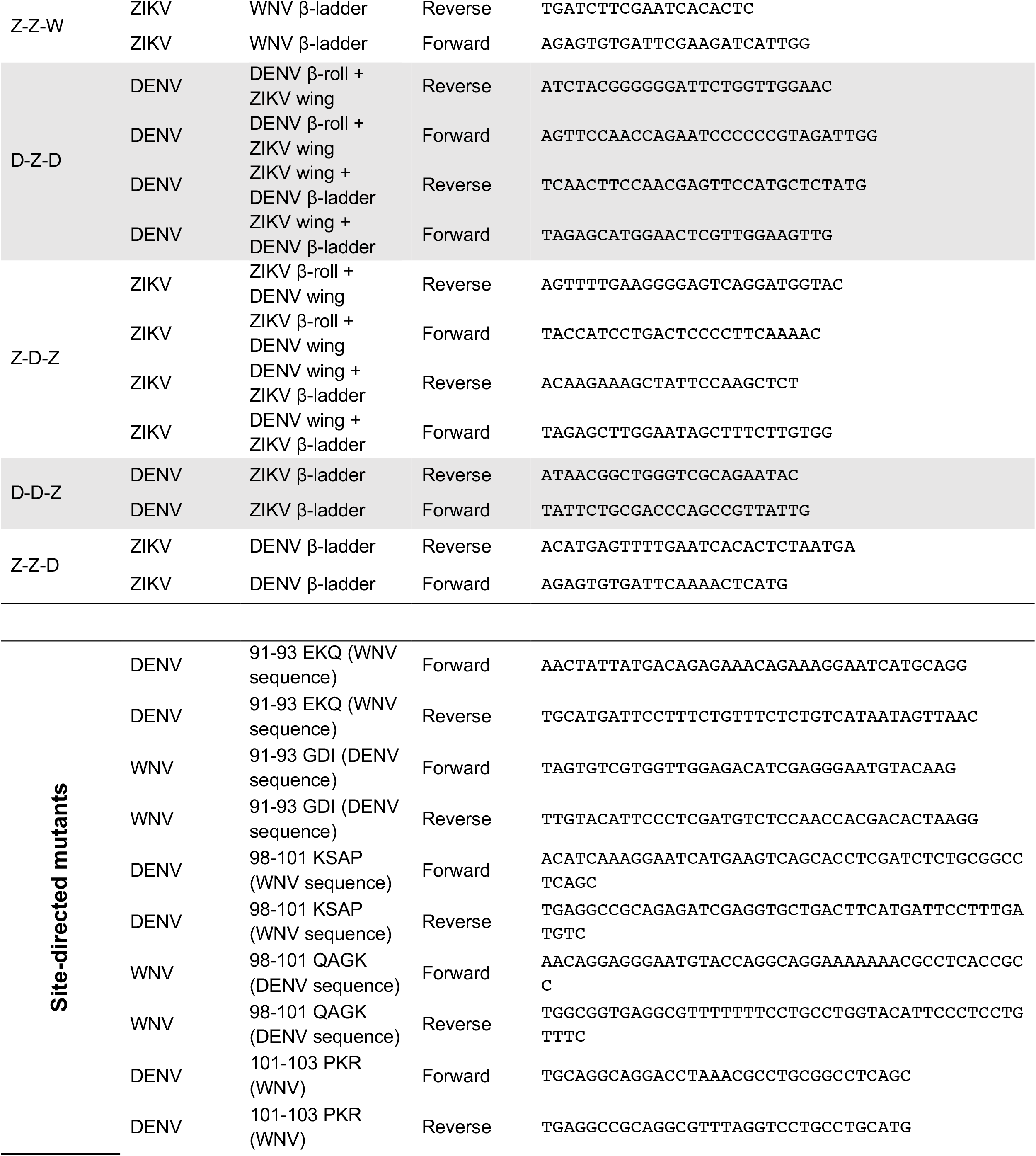
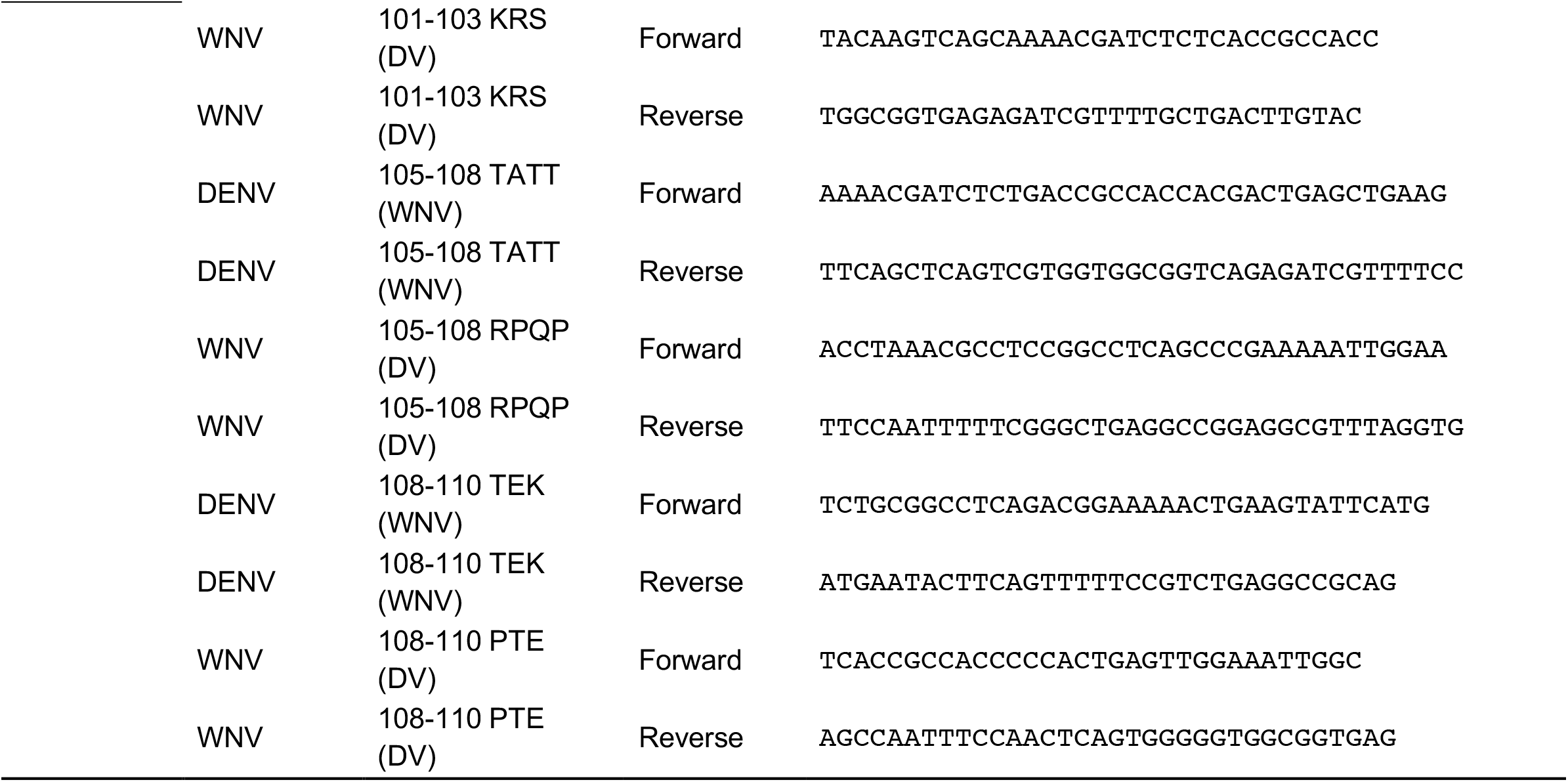
Primers used for mutagenesis in this study.

### NS1 protein production and purification

Plasmids containing WT or mutant NS1 sequence were transfected into FreeStyle 293F cells using polyethylenimine (PEI) (40K) (Sigma) according to the manufacturer’s instructions. 48 to 72 hours post-transfection, NS1-containing supernatants were collected, filtered through a 0.45 µm cellulose acetate membrane to remove cell debris, and stored at -80° prior to protein purification. The NS1-containing supernatants were thawed, mixed 1:1 with binding buffer (20 mM sodium phosphate, 500 mM sodium chloride, 20 mM imidazole, pH 7.4), and bound to HisPur cobalt resin (Thermo Fisher Scientific) with shaking for 2 hours at room temperature. The NS1-resin mixture then transferred to a column and washed 5 times in wash buffer (20 mM sodium phosphate, 500 mM sodium chloride, 25 mM imidazole, pH 7.4). NS1 was then eluted from the HisPur cobalt resin with elution buffer (20 mM sodium phosphate, 500 mM sodium chloride, 200 mM imidazole, pH 7.4) over 5 fractions. The purified NS1 stocks were then subjected to dialysis against 1X PBS for 48 hours at 4°C and concentrated using Amicon filters with 10,000 molecular weight cut-off (Millipore). The Pierce BCA protein quantitation kit (Thermo Fisher Scientific) was used to quantify the purified recombinant proteins according to manufacturer’s instructions. These proteins were used for all experiments within this study.

### SDS-PAGE and western blot

Recombinant proteins were collected in protein sample buffer (0.1 M Tris pH 6.8, 4% SDS, 4 mM EDTA, 286 mM 2-mercaptoethanol, 3.2 M glycerol, 0.05% bromophenol blue) and then resolved by SDS-PAGE. For native gels, the same protocol was followed except that the sample buffer contained no SDS and 2-mercaptoethanol. Proteins were then transferred onto nitrocellulose membranes and probed with primary antibodies diluted in Tris-buffered saline with 0.1% Tween20 (TBST) containing 5% nonfat dry milk. Membranes and antibodies were incubated overnight rocking at 4°C. The next day, membranes were washed three times with TBST before being probed with anti-mouse HRP secondary antibodies diluted in 5% milk in TBST at a dilution of 1:5,000 at room temperature for 1 hour. Afterwards, membranes were washed with TBST three more times before being developed with ECL reagents and imaged on a ChemiDoc system with Image Lab software (Bio-Rad). The following antibodies were used: mouse anti-His (MA1-21315, Thermo Scientific), goat anti-mouse HRP (405306, Biolegend).

### NS1 cell binding assays

To measure binding of WT and mutant NS1 proteins to HPMEC and HUVEC, 1×10^5^ cells were seeded on glass coverslips in 24-well plates. Cells were allowed to form a fully confluent monolayer for 3 days, with medium change every other day. On the day of experiment, 10 µg/mL (3 µg in 300 µL) of NS1 proteins were prepared in 10 µL medium, then added to the cells. Untreated wells were used as negative controls. NS1 and cells were incubated for 1 hour at 37°C. Mouse anti-6xHis antibody conjugated to Alexa Fluor 647 (Novus Biologicals) was then added at a dilution of 1:200, together with Hoechst 33342 (Immunochemistry) at a 1:2000 dilution for staining of nuclei, for 30 minutes at 37°C. Cells were then washed twice in 1X PBS followed by fixation in 4% formaldehyde diluted in 1X PBS (Thermo Fisher Scientific). Coverslips were mounted onto microscope slides on a drop of ProLong Gold (Thermo Fisher Scientific) and imaged using a Zeiss LSM 710 inverted confocal microscope (CRL Molecular Imaging Center, UC Berkeley). Images were processed using ImageJ software.

### Trans-endothelial electrical resistance (TEER)

The trans-endothelial electrical resistance assay was used to measure the functional effect of NS1 on endothelial barrier function in HPMEC as previously described (33). Briefly, 1×10^5^ cells (HPMEC) were seeded in 300 µL of medium on the polycarbonate membrane insert of a trans-well (Transwell permeable support, 0.4 µm, 6.5 mm insert; Corning Inc.). The trans-well is placed in a well on a 24-well plate, becoming the apical (upper) chamber. 1.5 mL of media is added to the basolateral (lower) chamber. Cells were allowed to form a monolayer for 3 days with media changes in both apical and basolateral chambers every day, until the inter-chamber electrical resistance reaches about 60 Ω difference between trans-wells seeded with (∼150 Ω) and without cells (∼90 Ω). On the day of experiment, 2.5 or 5 µg/mL of indicated NS1 proteins (0.75 or 1.5 µg proteins, respectively) were mixed with media up to 10µL, and added to the apical chambers of the trans-wells. Electrical resistance between the apical and basolateral chambers is measured in ohms using an Epithelial Volt Ohm Meter (EVOM) with an electrode pair (World Precision Instruments), at the times indicated in the figures. Trans-wells containing no cells and untreated trans-wells containing only cells were used as negative controls to calculate the baseline electrical resistance at each timepoint. Relative TEER is calculated as a ratio of resistance values ((Ω_experimental_ – Ω_media only_) / (Ω_untreated cells_ – Ω_media only_)). For area under the curve (AUC) analyses, the net AUC was taken from all curves using baseline of Y=1.

### Mouse model of localized vascular leak

Five-to eight-week-old WT C57BL/6 male mice were purchased from the Jackson Laboratory (Bar Harbor, ME) and maintained under specific pathogen-free conditions at the University of California, Berkeley, Animal Facility. Mice were housed in a controlled temperature environment on a 12-hour light/dark cycle, with food and water provided *ad libitum*. All experimental procedures involving animals were pre-approved by the Animal Care and Use Committee (ACUC) of the University of California, Berkeley. Three to four days prior to experiment, the dorsal dermises of 6-to 10-week-old WT C57BL/6 female mice (Jackson Laboratory) were shaved using hair clippers, and residual hair removed using Nair (Church & Dwight). On the day of experiment, 15 µg of WT or mutant NS1 was mixed with PBS in a total volume 50 µL each. NS1 mixtures and PBS were then injected intradermally (ID) into discrete spots in the shaved mouse dermis. Immediately following ID injections, 25 µg of 10-kDa dextran conjugated to Alexa Fluor 680 (1 mg/mL; Sigma) was delivered intravenously (IV) through the retro-orbital route. Two hours post-injection, mice were euthanized, and the dorsal dermis was removed and placed in Petri dishes. The dermis was then placed on a fluorescent scanner (LI-COR Odyssey CLx Imaging System) to visualize the fluorescence signal accumulation at a wavelength of 700 nm. Vascular leak at the ID injection sites was quantified using Image Studio software (LI-COR Biosciences) as described previously (30, 31).

### Statistics

All quantitative analyses were conducted, and all data were plotted, using GraphPad Prism 9 software. Experiments were repeated at least 3 times, to ensure reproducibility. All experiments were designed and performed with both positive and negative controls (indicated in the figures), which were used for inclusion/exclusion determination. For immunofluorescence microscopy experiments, images of random fields were captured. For all experiments with quantitative analysis, data are displayed as mean ± standard error of the mean (SEM). All cell binding and TEER quantitative data were analyzed using a One-way ANOVA analysis with Tukey’s multiple comparisons test. For the localized dermal leak experiments, a non-parametric, unpaired Mann-Whitney U test was used to determine statistical significance between groups. The resulting p-values from the above statistical tests were displayed as n.s., not significant; p >0.05; *p <0.05; **p <0.01; ***p <0.001; ****p <0.0001.

